# Loss of Spindly sensitizes EML4-ALK v3 lung cancer cells to HSP90 inhibitors

**DOI:** 10.1101/2022.06.08.495301

**Authors:** Marco P. Licciardello, Chi Zhang, Anh T. Le, Robert C. Doebele, Paul A. Clarke, Paul Workman

## Abstract

Heat Shock Protein 90 inhibitors (HSP90i) have shown activity in EML4-ALK^+^ non-small cell lung cancer (NSCLC) patients but clinical responses have been heterogeneous. It has been suggested that distinct EML4-ALK variants may have a differential impact on the response to HSP90 inhibition. Here, we show that NSCLC cells harboring the most common EML4-ALK variant 1 (v1) or variant 3 (v3) are in fact similarly sensitive to HSP90i. To discover new genetic alterations that could be involved in stratifying sensitivity, we performed a genome-wide CRISPR/Cas9 knockout screen and found that loss of Spindly increases the sensitivity of EML4-ALK v3, but not v1, NSCLC cells to low concentrations of HSP90i from three distinct chemical families. Upon loss of Spindly, prolonged exposure to low concentrations of HSP90i impairs chromosome congression and cellular fitness. Collectively, our data suggest that mutations leading to loss of Spindly in EML4-ALK v3 NSCLC patients may increase sensitivity to low doses of HSP90i.

## Introduction

Between 3-7% of NSCLC patients harbor the fusion oncogene *EML4-ALK*. Echinoderm microtubule-associated protein-like 4 (EML4) is a microtubule-stabilizing protein involved in the formation of the mitotic spindle (1; 2). Anaplastic lymphoma kinase (ALK) is a transmembrane tyrosine kinase possibly playing a role in fetal brain development and otherwise poorly expressed in adult tissues (3). At least 15 EML4-ALK variants have been reported, each carrying the intracellular kinase domain of ALK at the C-terminal fused to different portions of EML4 at the N-terminal. EML4-ALK v1 and EML4-ALK v3 are the most common variants and account for the majority of all cases (4). The longer v1 localizes to cytoplasmic granules and retains a portion of the tandem atypical propeller in EMLs (TAPE) domain of EML4, which confers structural instability; in contrast, the shorter v3 does not retain portions of the TAPE domain, is more stable and localizes to both cytoplasmic granules and microtubules (5; 6; 7). All variants retain the trimerization domain of EML4, which leads to the cancer-addicting constitutive activation of ALK and downstream MAPK/PI3K signaling. NSCLC patients harboring the fusion oncogene *EML4-ALK* have been successfully treated with ALK tyrosine kinase inhibitors such as crizotinib or, more recently, alectinib and lorlatinib; however, patients inevitably develop resistance to these targeted therapies and require alternative treatments (8; 9).

The molecular chaperone HSP90 is one of the major players of the proteostasis network ensuring correct folding of protein clients in physiological conditions and in response to stress (10). HSP90 also plays an important role in cancer development as many oncogenes rely on its adenosine triphosphate (ATP)-driven chaperoning activity for their stability and function (11). For this reason, HSP90 and other components of the proteostasis network have been investigated as potential targets for the treatment of cancer (12). All clinical stage HSP90i target the ATP-binding pocket in the N-terminal domain of the chaperone (13; 14). These small molecules are usually administered at doses that lead to client misfolding followed by proteasomal degradation and induction of heat shock response genes including *HSPA1A* (HSP72) and *HSPB1* (HSP27) (15).

EML4-ALK v1 has been shown to interact with HSP90 and to be rapidly depleted in NSCLC cells exposed to HSP90i (16; 17). These observations have led to the inclusion of EML4-ALK^+^ NSCLC patients in clinical trials of HSP90i. Interestingly, HSP90i such as the geldanamycin-based retaspimycin (IPI-504) and the resorcinol-based ganetespib (STA-9090) and luminespib (AUY922) have shown activity in NSCLC patients harboring the *EML4-ALK* fusion oncogene. However, responses have been heterogeneous and, despite acceptable tolerability, patients have experienced adverse events at the recommended doses (18; 19; 20). It has been hypothesized that the heterogeneous responses to HSP90i observed in the clinic might have been the result of a molecular stratification of EML4-ALK^+^ NSCLC patients that has so far ignored variant-specific structural features (21). Indeed, endogenous EML4-ALK v3 was not depleted in NSCLC cells upon treatment with ganetespib in contrast to EML4-ALK v1 (22); however, the *in vitro* exposure was extended for only 6 h. Moreover, while the exogenous expression of EML4-ALK variants in Ba/F3 cells led to a difference in sensitivity to ganetespib after 24 h, viability measurements at longer exposures and in NSCLC cells expressing endogenous EML4-ALK variants were not included (22). In another study, no substantial difference in IC_50_ was observed in concentration-response curves when NSCLC cells harboring either v1 or v3 were exposed to ganetespib for 72 h (23). Clearly, the contribution of different EML4-ALK variants to the sensitivity of NSCLC cells to HSP90i requires further elucidation. Here, we confirm that v1 is depleted more rapidly than v3 upon exposure of NSCLC cells to HSP90i. However, NSCLC cells harboring different EML4-ALK variants show little or no difference in sensitivity upon treatment with HSP90i for 72 h or longer.

Heat shock proteins facilitate the folding of mutated protein clients and thus function as capacitors buffering genetic variations (24) and displaying many potential synthetic lethal interactions (25). To uncover such synthetic lethal partners and further characterize the mechanism of action of HSP90i in the context of EML4-ALK^+^ NSCLC, we performed a genome-wide clustered regularly interspaced palindromic repeats (CRISPR)/Cas9 knockout screen in NSCLC cells harboring either v1 or v3 and exposed to a non-lethal concentration of an HSP90i. We discovered that the knockout of *SPDL1*, the gene encoding the protein Spindly, increases the sensitivity of NSCLC cells harboring EML4-ALK v3, but not v1, to low concentrations of structurally distinct HSP90i. We show that, upon loss of Spindly, the prolonged exposure of v3 cells to low concentrations of HSP90i leads to impairment of chromosome congression and cellular fitness. Overall, our findings suggest that upon loss or depletion of Spindly, EML4-ALK v3 NSCLC patients might be more responsive to low doses of HSP90i and less likely to experience the adverse events observed in past clinical trials.

## Results

### EML4-ALK^+^ NSCLC cells show similar sensitivity to HSP90i

Previous reports have shown that, in contrast to v3, EML4-ALK v1 is rapidly depleted in NSCLC cells exposed to HSP90i (17; 22). We aimed to reproduce and further investigate these results using HSP90i from three distinct chemical families: the geldanamycin-derived tanespimycin (17-*N*-allylamino-17-demethoxygeldanamycin, 17-AAG), the resorcinol-based luminespib and the purine-scaffold BIIB021 (**Figure 1A**). Indeed, we observed a concentration-dependent depletion of EML4-ALK v1 in H3122 NSCLC cells exposed to tanespimycin or luminespib for 24 h whereas, in contrast, EML4-ALK v3 was only partly depleted at the highest concentrations in H2228 NSCLC cells (**Figures 1B** and **S1A**).

**Figure 1.**
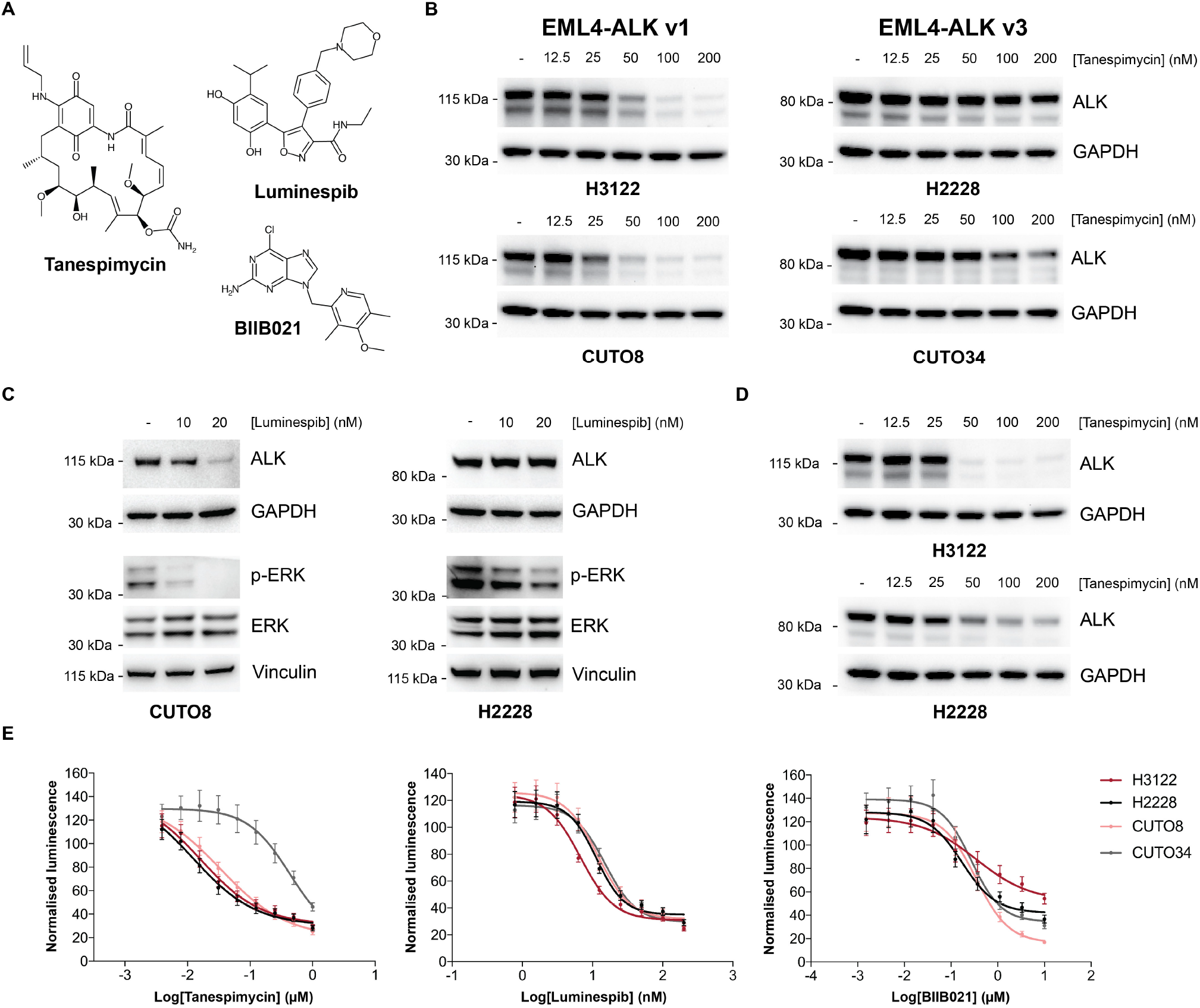
EML4-ALK^+^ NSCLC cells are equally sensitive to HSP90i. (**A**) Structures of HSP90i from different chemical families. (**B**) Western blotting of EML4-ALK^+^ NSCLC cells exposed to tanespimycin at the indicated concentrations for 24 h. GAPDH was used as a loading control. (**C**) Western blotting of CUTO8 and H2228 cells exposed to luminespib at the indicated concentrations for 24 h. GAPDH and Vinculin were used as loading controls. (**D**) Western blotting of H3122 and H2228 cells exposed to tanespimycin at the indicated concentrations for 72 h. GAPDH was used as a loading control. (**E**) Concentration-response curves of EML4-ALK^+^ NSCLC cells exposed to different HSP90i as indicated for 72 h. Data are normalized to vehicle-treated cells set to 100. Error bars, s.d. of 6 biological replicates.

The availability of EML4-ALK^+^ NSCLC cell lines is very limited and most studies on this specific type of lung cancer are usually performed using only H3122 and H2228 cells. To corroborate our results, we sourced two additional patient-derived EML4-ALK^+^ NSCLC cell lines, namely CUTO8 (*EML4-ALK* E13;A20, v1) and CUTO34 (*EML4-ALK* E6;A20, v3) cells. Akin to our observations in H3122 and H2228 cells, EML4-ALK v1 was depleted in a concentration-dependent manner in CUTO8 cells whereas, in contrast, a moderate depletion of EML4-ALK v3 was detected in CUTO34 cells only at higher concentrations upon exposure to HSP90i for 24 h (**Figures 1B** and **S1A**). Depletion of EML4-ALK v1 in CUTO8 cells upon treatment with luminespib for 24 h led to decreased MAPK signaling, which is triggered downstream of constitutive ALK activity (**Figure 1C**). Interestingly, we observed reduced MAPK signaling after 24 h also in H2228 cells exposed to concentrations of luminespib that did not lead to depletion of EML4-ALK v3 (**Figure 1C**). Of note, we detected further depletion of EML4-ALK v3 in H2228 cells upon exposure to tanespimycin for 72 h (**Figure 1D**). Whereas treatment with luminespib at concentrations leading to decreased MAPK signaling did not affect the cellular fitness of CUTO8 and H2228 cells at 24 h (**Figure S1B**), we observed loss of cellular fitness but no substantial difference in sensitivity when EML4-ALK^+^ NSCLC cells were treated with the 3 structurally distinct HSP90i for 72 h (**Figure 1E**). CUTO34 cells are more resistant to tanespimycin but this is explained, at least in part, by the very low levels of NAD(P)H quinone dehydrogenase 1 (NQO1), a flavoenzyme oxidoreductase that metabolically activates tanespimycin (26; 27), expressed in these cells (**Figure S1C**). Exposure to tanespimycin, luminespib or BIIB021 for 72 h induced apoptosis at similar concentrations in all EML4-ALK^+^ NSCLC cells as indicated by the increased amount of cleaved PARP (**Figure S1D**). Additionally, we assessed the cellular fitness of EML4-ALK^+^ NSCLC cells exposed to different HSP90i for 12 d. EML4-ALK v1 H3122 cells, but not CUTO8 cells, were slightly more sensitive to HSP90i (**Figure S1E**). Overall, these results confirmed that EML4-ALK v1 is depleted more rapidly although v3 is also eventually depleted, at least in part, upon perturbation of the proteostasis network in NSCLC cells. Exposure of EML4-ALK^+^ NSCLC cells to HSP90i leads to decreased MAPK signaling even before client depletion and regardless of the EML4-ALK variant expressed. The depletion of EML4-ALK and the decrease of downstream cancer-addicting MAPK signaling explain the sensitivity of EML4-ALK^+^ NSCLC cells to inhibition of HSP90. As we did not observe any substantial difference in sensitivity to HSP90i among our EML4-ALK^+^ NSCLC cells, we argue that the expression of different EML4-ALK variants alone is unlikely to explain the heterogeneous responses observed in EML4-ALK^+^ NSCLC patients treated with HSP90i.

### Knockout of *SPDL1* sensitizes v3 cells to low concentrations of HSP90i

To discover additional genes that could be involved in stratifying the sensitivity to HSP90i, we introduced the Cas9 endonuclease from *S. pyogenes* in EML4-ALK v1 H3122 and EML4-ALK v3 H2228 cells via lentiviral transduction. We confirmed sufficient Cas9 protein levels using western blotting and activity of the endonuclease by means of a flow cytometry-based assay (**Figures S2A** and **S2B**). We then expanded the CRISPR single guide RNA (sgRNA) Brunello library (28) (**Figure S2C**) and used it to perform a pooled genome-wide knockout screen on Cas9-expressing H3122 and H2228 cells (**Figure 2A**). Cells transduced with the Brunello library were exposed to either DMSO or 12 nM tanespimycin for 12 d to capture cell cycle-and epigenetic-dependent effects. At this low concentration, tanespimycin induces HSP72, a marker of HSP90 inhibition, but does not deplete EML4-ALK (**Figure S2D**). We collected genomic DNA samples and assessed the distribution of sgRNAs before (T_0_) and after drug treatment (T_12_) using next-generation sequencing.

**Figure 2.**
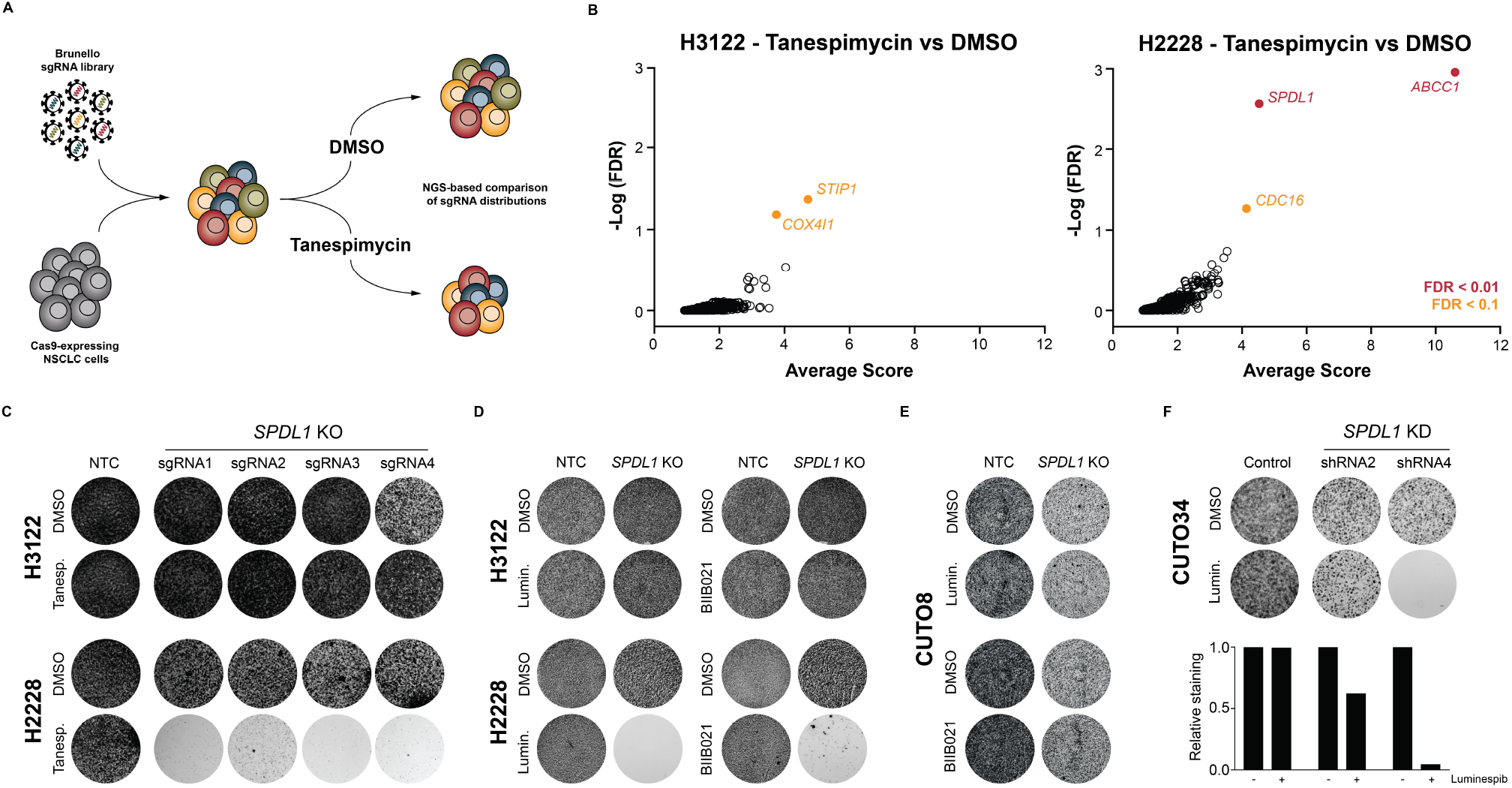
Loss of Spindly is synthetic lethal with low concentrations of HSP90i in EML4-ALK v3 NSCLC cells. (**A**) Schematic representation of the genome-wide CRISPR/Cas9 pooled knockout screen. (**B**) The average score and the -Log of the FDR are plotted for EML4-ALK v1 H3122 (left) and EML4-ALK v3 H2228 (right) cells. Each circle represents a gene. Hits with a FDR<0.1 are indicated in orange, hits with a FDR<0.01 are indicated in red. (**C**) Crystal violet staining of NTC and *SPDL1* KO H3122 and H2228 cells exposed to DMSO or 12 nM tanespimycin for 12 d. (**D**) Crystal violet staining of NTC and *SPDL1* KO H3122 and H2228 cells exposed to DMSO or 6 nM luminespib or 20 nM BIIB021 for 12 d. (**E**) Crystal violet staining of NTC and *SPDL1* KO CUTO8 cells exposed to DMSO or 6 nM luminespib or 30 nM BIIB021 for 12 d. (**F**) Crystal violet staining and quantification of control and *SPDL1* KD CUTO34 cells exposed to DMSO or 6 nM luminespib for 12 d.

Biological replicates showed good correlation (**Figure S2E**) and the sgRNA ratio between the T_12_ DMSO sample and the T_0_ sample did not change for the vast majority of the 1,000 non-targeting control (NTC) sgRNAs included in the Brunello library, as expected (**Figure S2F**). We compared sgRNA distributions using the STARS algorithm (28) to rank genes. Overall, 1017 genes were depleted (FDR<0.1) in the T_12_ DMSO sample versus the T_0_ sample of the H2228 screen (**Figure S2G**; **Table S1**). These genes decrease the cellular fitness of H2228 cells when knocked out regardless of treatment with tanespimycin. We compared this list with a catalog of 553 pan-cancer core essential genes that has been recently described (29) and found an overlap of 74%, which increased to 84% when we relaxed the significance threshold (**Figure S2G**). We observed a slightly smaller number of depleted genes when we compared the T_12_ DMSO sample with the T_0_ sample of the H3122 screen and an overlap of up to 64% with the pan-cancer core essential genes catalog (**Figure S2G**; **Table S1**). Overall, these results demonstrate the technical success of the CRISPR/Cas9 screen.

Interestingly, top-scoring genes depleted in the tanespimycin T_12_ sample versus the DMSO T_12_ sample— that is genes that increase the sensitivity to tanespimycin when knocked out — were different for H3122 and H2228 cells. In EML4-ALK v1 H3122 cells, the top-scoring depleted genes (FDR<0.1) were *COX4I1*, which encodes a subunit of complex IV of the mitochondrial respiratory chain, and *STIP1* (**Figure 2B**; **Table S2**). *STIP1* encodes the co-chaperone HOP, which is involved in the transferring of client proteins between the HSP70 and the HSP90 chaperone machineries (30). The structural instability of EML4-ALK v1 likely explains the increased sensitivity of H3122 cells to tanespimycin upon further perturbation of the proteostasis network following knockout of *STIP1*. In EML4-ALK v3 H2228 cells, the top-scoring depleted genes were *CDC16* (FDR<0.1), which encodes a component of the anaphase promoting complex, and the high-confidence *SPDL1* and *ABCC1* (FDR<0.01) (**Figure 2B**; **Table S2**). Notably, *ABCC1* encodes the multidrug resistance protein 1 (MRP1), which has been shown to be involved in resistance to geldanamycin analogues (31). Loss of MRP1 increased the sensitivity of H2228 cells to the geldanamycin-derived HSP90i tanespimycin in single sgRNA experiments, validating our screening approach (**Figure S3A**). Knockout of *ABCC1* did not increase the sensitivity of H3122 cells to tanespimycin, most likely because of lower expression in this cell line (**Figure S3B**).

*SPDL1*, which encodes the protein Spindly, scored as the second high-confidence top gene in EML4-ALK v3 H2228 cells but its loss did not increase the sensitivity to tanespimycin in EML4-ALK v1 H3122 cells (**Figure 2B**). Spindly recruits the motor complex dynein to kinetochores and is involved in the formation of the mitotic spindle and chromosome congression (32; 33). Upon entering prometaphase, kinetochores assemble on the centromeres of sister chromatids and form attachments with microtubules. A complex quality control system ensures that bi-orientated amphitelic kinetochore-microtubule (k-MT) attachments are correctly established and that chromosomes congress to the metaphase plate to release the spindle assembly checkpoint (SAC) and promote entry into anaphase (34). Dynein mediates the movement of laterally attached chromosomes to the spindle poles facilitating their congression to the metaphase plate where they can form stable bi-orientated k-MT attachments (35). Hence, loss of Spindly may lead to impairment of chromosome congression as dynein cannot be efficiently recruited to kinetochores in the absence of Spindly. Indeed, depletion of Spindly by RNA interference has been reported to induce mitotic delay and chromosome congression defects in human cells (36). However, alternative mechanisms can ensure satisfactory chromosome segregation in the absence of dynein (35).

In contrast to v1, EML4-ALK v3 localizes to microtubules (5; 6) and might be associated, directly or indirectly, with other proteins involved in the formation of the mitotic spindle. For this reason and because of the high-confidence score of the *SPDL1* gene in v3 H2228 cells, we further investigated the role played by depletion of Spindly in increasing the sensitivity of EML4-ALK v3 NSCLC cells to low concentrations of HSP90i. We validated the results of the screen transducing H3122 and H2228 cells with a single NTC sgRNA (hereinafter referred to as NTC cells) or single sgRNAs targeting *SPDL1* (hereinafter referred to as *SPDL1* KO cells) (**Figure S3C**) and confirmed that loss of Spindly increases the sensitivity to low concentrations of tanespimycin only in v3 H2228 cells (**Figure 2C**). Of note, we observed a similar increase in sensitivity to low concentrations of luminespib and BIIB021 upon knockout of *SPDL1* in EML4-ALK v3 H2228 cells but not in EML4-ALK v1 H3122 and CUTO8 cells (**Figures 2D, 2E** and **S3D**). Moreover, in contrast to EML4-ALK v1 CUTO8 cells, EML4-ALK v3 CUTO34 cells showed increased sensitivity to low concentration of HSP90i upon knock-down of Spindly using two short hairpin RNA (shRNA) constructs (**Figures 2F, S3E** and **S3F**). Previous reports have described a synergistic interaction between HSP90i and microtubule-targeting agents in selected NSCLC cell lines (37; 38). Of note, EML4-ALK v3 H2228 cells are very sensitive to taxanes and the addition of ganetespib does not lead to a further increase of antitumor activity in NSCLC xenograft models (39). Exposure to the highest concentration of tanespimycin that is synthetic lethal with loss of Spindly did not increase the sensitivity of EML4-ALK v3 H2228 cells to non-lethal concentrations of paclitaxel (**Figure S3G**). Moreover, our CRISPR/Cas9 screen did not reveal a general increase in sensitivity to low concentrations of HSP90i upon depletion of proteins involved in spindle or kinetochore assembly. Overall, these data indicate that the specific loss of Spindly increases the sensitivity of NSCLC cells harboring EML4-ALK v3, but not v1, to low concentrations of structurally distinct HSP90i.

### *SPDL1* KO v3 cells treated with HSP90i accumulate in mitosis

EML4-ALK v3 may interact with proteins involved in the formation of the mitotic spindle, which in turn could affect its stability. We therefore asked whether loss of Spindly increases sensitivity to HSP90i by decreasing the stability of EML4-ALK v3 in NSCLC cells. We exposed NTC and *SPDL1* KO H2228 cells to different concentration of tanespimycin for 24 and 72 h and observed no change in the stability of EML4-ALK v3 (**Figure S4A**). Interestingly, we detected no increase in sensitivity when we compared *SPDL1* KO to NTC H2228 cells in a concentration-response treatment with tanespimycin over 72 h (**Figure S4B**). However, we observed growth impairment of *SPDL1* KO H2228, but not NTC H2228 cells, upon exposure to a low concentration of tanespimycin for 10 d or longer (**Figure 3A**). Notably, knock-out of *SPDL1* alone had only a minimal effect on the growth of H2228 cells (**Figure 3A**), supporting Spindly as a potential synthetic lethal partner of HSP90i in the context of EML4-ALK v3 NSCLC.

**Figure 3.**
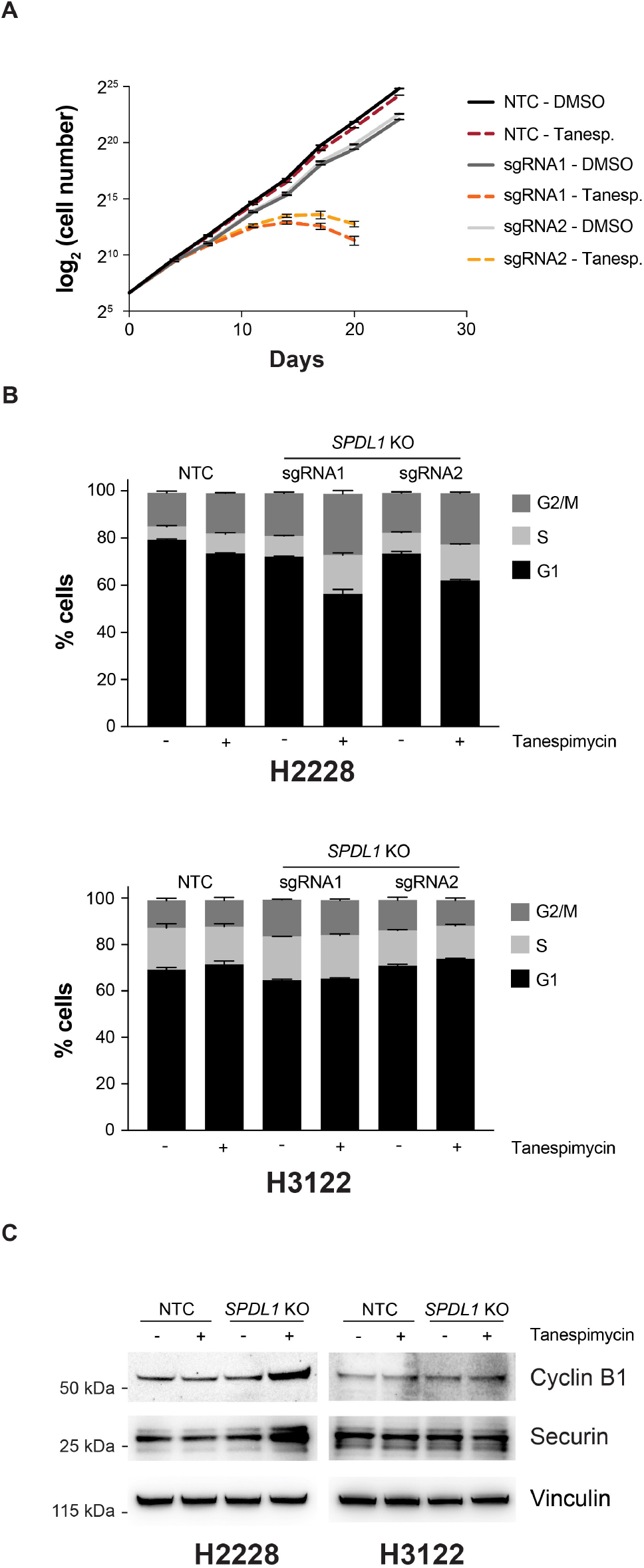
Knockout of *SPDL1* and low concentrations of HSP90i lead to EML4-ALK v3 NSCLC cells accumulation in mitosis. **(A)** Growth curve of NTC and *SPDL1* KO H2228 cells exposed to DMSO or 12 nM tanespimycin for the indicated time. Error bars, s.d. of 3 biological replicates. **(B)** Cell cycle analysis by flow cytometry of NTC and *SPDL1* KO H2228 (top) and H3122 (bottom) cells exposed to DMSO or 6 nM tanespimycin for 10 d. Error bars, s.d. of 3 biological replicates. (**C**) Western blotting of NTC and *SPDL1* KO H2228 and H3122 cells treated as in panel (B). Vinculin was used as a loading control.

As Spindly is involved in the formation of the mitotic spindle and chromosome congression, we determined effects on the cell cycle. The exposure of NTC H2228 cells to a low concentration of tanespimycin for 10 d or the knockout of *SPDL1* alone led to a small increase in the number of cells in S and G2/M of around 3% (**Figures 3B** and **S4C**). However, upon exposure of *SPDL1* KO H2228 cells to tanespimycin, this increase was around 10% when compared to untreated NTC H2228 cells (**Figures 3B** and **S4C**). We observed a small increase in the number of cells in G2/M of around 3% also upon knockout of *SPDL1* alone in H3122 cells. However, in contrast to *SPDL1* KO EML4-ALK v3 H2228 cells, there was no further increase when *SPDL1* KO EML4-ALK v1 H3122 cells were exposed to tanespimycin for 10 d (**Figures 3B** and **S4C**). Moreover, we detected increased levels of the prometaphase/metaphase markers cyclin B1 and securin in *SPDL1* KO v3 H2228 cells, but not in *SPDL1* KO v1 H3122 and CUTO8 cells, exposed to low concentrations of HSP90i for 10 d (**Figures 3C** and **S4D**). This suggests that EML4-ALK v3, but not v1, NSCLC cells accumulate in prometaphase/metaphase upon concomitant loss of Spindly and prolonged treatment with low concentrations of HSP90i.

### Low concentrations of HSP90i induce congression defects upon loss of Spindly

We examined spindle formation and chromosome congression via immunofluorescence. Both NTC and *SPDL1* KO EML4-ALK v3 H2228 cells showed around 9-10% of mitotic cells with multipolar spindles and this value did not change considerably upon exposure to a low concentration of tanespimycin for 10 d (**Figure S4E**). The vast majority of mitotic NTC H2228 cells showed no chromosome congression impairment and the percentage of cells with a disrupted spindle increased only by around 5% upon prolonged exposure to tanespimycin and by around 10% upon knockout of *SPDL1* alone (**Figure 4A**). In contrast, we observed an increase of around 35% in the percentage of cells with uncongressed chromosomes and a severely disrupted spindle when *SPDL1* KO H2228 cells were exposed to tanespimycin for 10 d (**Figure 4A**). Additionally, the percentage of NTC H2228 cells with micronuclei increased by around 9% upon exposure to tanespimycin and we observed a similar increase upon knockout of *SPDL1* alone in H2228 cells (**Figure 4B**). Of note, prolonged treatment of *SPDL1* KO H2228 cells with tanespimycin led to a 20% increase in the percentage of cells with micronuclei (**Figure 4B**). Interestingly, we detected an increase in intact cells with fragmented nuclei of around 14% when we exposed NTC H2228 cells to a low concentration of tanespimycin for 10 d. Strikingly, while loss of Spindly alone did not lead to the same observation, the percentage of cells with fragmented nuclei increased by around 28% when *SPDL1* KO H2228 cells were treated with tanespimycin (**Figure 4C**). Overall, these results suggest that while prolonged exposure to a low concentration of HSP90i or loss of Spindly alone might in part affect cell cycle progression and chromosome segregation without substantially interfering with cellular fitness, the concomitant presence of these two perturbations leads to greater impairment of chromosome congression and, after a number of cell divisions, growth inhibition in EML4-ALK v3 NSCLC cells.

**Figure 4.**
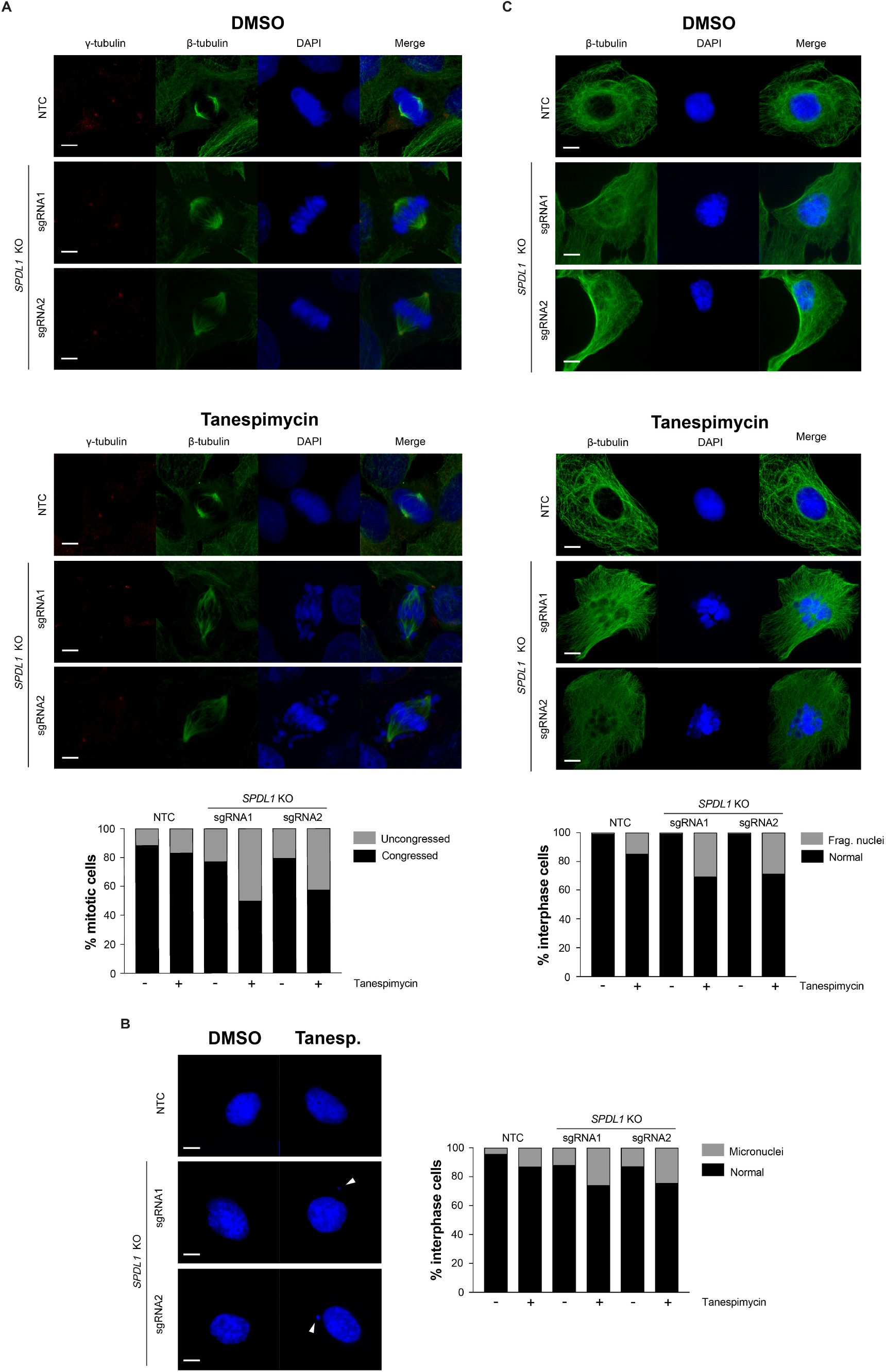
Loss of Spindly induces chromosome congression impairment in EML4-ALK v3 NSCLC cells upon treatment with HSP90i. (**A-C**) Immunofluorescence analysis and quantification of NTC and *SPDL1* KO H2228 cells exposed to DMSO or 6 nM tanespimycin for 10 d. For the analysis of chromosome congression, more than 200 mitotic cells per condition were counted to determine percentages. Bipolar spindles with clear chromosome congression defects as well as multipolar spindles were counted as ‘uncongressed’. For the analysis of micronuclei and fragmented nuclei, more than 400 interphase cells per condition were counted to determine percentages. Scale bar, 5 µm.

### Increased chromosomal passenger complex activity reduces cellular fitness

To further elucidate the mechanism behind this synthetic lethal interaction, we performed gene expression profiling by RNA sequencing of NTC and *SPDL1* KO EML4-ALK v3 H2228 cells exposed to either DMSO or a low concentration of tanespimycin for 10 d. As we observe growth impairment after prolonged treatment, we aimed to explore the transcriptional landscape of NTC and *SPDL1* KO H2228 cells at a later time-point rather than focusing on the immediate response of these cells to HSP90i. Biological replicates showed excellent correlation (**Figure S5A**) and different sgRNAs targeting *SPDL1* produced very similar results supporting on-target effects (**Figure 5A**). Overall, loss of Spindly significantly perturbed the expression of less than 100 genes (fold change>2, FDR<0.01) while prolonged treatment of NTC H2228 cells with a low concentration of tanespimycin significantly altered the expression of just under 600 genes including members of the heat shock response (**Figure S5B**; **Table S3**). However, the concomitant presence of these two perturbations induced significant changes in the expression of more than 2000 genes (**Figure 5A**; **Table S3**).

**Figure 5.**
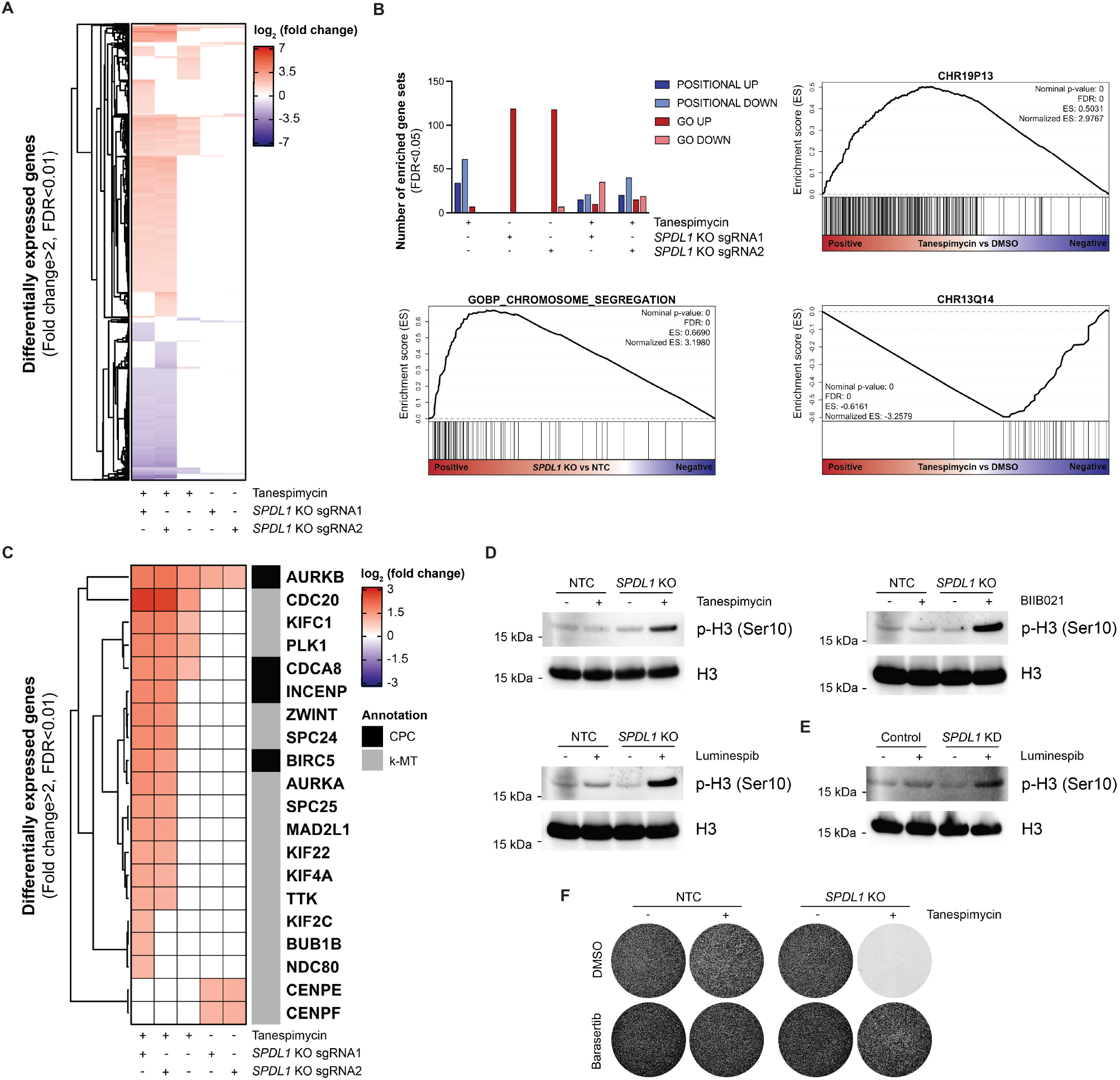
Low concentrations of HSP90i increase CPC activity at kinetochores upon loss of Spindly. (**A**) Heatmap showing gene expression changes in NTC and *SPDL1* KO H2228 cells exposed to DMSO or 6 nM tanespimycin for 10 d. The fold change to the corresponding NTC or DMSO control is illustrated. Only fold changes>2 with FDR<0.01 were considered. (**B**) Quantification and representative examples of gene sets significantly enriched by GSEA among genes up-or downregulated in NTC and *SPDL1* KO H2228 treated as in panel (A). Only gene sets enriched with FDR<0.05 were considered. (**C**) Heatmap showing genes involved in the formation of k-MT attachments differentially expressed in NTC and *SPDL1* KO H2228 treated as in panel (A). The fold change to the corresponding NTC or DMSO control is illustrated. Only fold changes>2 with FDR<0.01 were considered. (**D**) Western blotting of NTC and *SPDL1* KO H2228 cells exposed to DMSO or 6 nM tanespimycin or 3 nM luminespib or 10 nM BIIB021 for 10 d. Histone H3 was used as a loading control. (**E**) Western blotting of control and *SPDL1* KD CUTO34 cells exposed to DMSO or 3 nM luminespib for 10 d. Histone H3 was used as a loading control. (**F**) Crystal violet staining of NTC and *SPDL1* KO H2228 cells exposed to DMSO or 12 nM tanespimycin for 12 d with or without the Aurora B inhibitor barasertib (2.5 nM).

Interestingly, gene set enrichment analysis (GSEA) (40) revealed enrichment of 95 positional gene sets (FDR<0.05) upon prolonged exposure of H2228 cells to a low concentration of tanespimycin alone (**Figure 5B**; **Table S4**), suggesting concordant changes in the expression levels of neighboring genes on chromosomes (41). We observed significant enrichment (FDR<0.05) of gene ontology signatures associated with chromosome congression among genes upregulated upon knockout of *SPDL1* alone (**Figure 5B**; **Table S4**). GSEA revealed both positional and gene ontology gene sets associated with chromosome congression in *SPDL1* KO H2228 cells exposed to a low concentration of tanespimycin, although fewer gene sets scored at FDR<0.05 in these conditions (**Figure 5B**; **Table S4**).

We then focused on genes encoding proteins that are known to play a role in the formation of k-MT attachments and chromosome congression. Upon knockout of *SPDL1* in H2228 cells, we observed an increase in the expression of *CENPE, CENPF* and *AURKB* (**Figure 5C**; **Table S3**). CENPF contributes to dynein recruitment (42) while CENPE is involved in alternative chromosome movements (35) that might offset the depletion of dynein at kinetochores following loss of Spindly. *AURKB* encodes Aurora B kinase, a subunit of the chromosomal passenger complex (CPC) together with INCENP, survivin (*BIRC5*) and borealin (*CDCA8*). This complex plays a crucial role in the formation of k-MT attachments mainly via the kinase activity of Aurora B (43). One of the main substrates of the CPC is Ser10 of histone H3 at centromeres and the phosphorylation of this amino acid by Aurora B contributes to the recruitment of the first elements of the kinetochore complex; the CPC phosphorylates additional substrates at kinetochores and these post-translational modifications decrease the stability of k-MT attachments (44). When erroneous k-MT attachments are generated, the CPC destabilizes these faulty connections and facilitates correction to bi-orientated amphitelic k-MT attachments, which are then stabilized by the dephosphorylation of Aurora B substrates at kinetochores. Importantly, all subunits of the CPC are equally necessary for the kinase activity of Aurora B at kinetochores (43).

The exposure of EML4-ALK v3 NTC H2228 cells to tanespimycin induced the expression of 2 genes encoding subunits of the CPC, namely *AURKB* and *CDCA8*, together with other genes encoding proteins involved in the formation of k-MT attachments such as *KIFC1* (HSET) and *PLK1* (**Figure 5C**). We observed induced expression of a greater number of genes whose protein products are involved in the formation of k-MT attachments, including all subunits of the CPC, upon prolonged exposure of *SPDL1* KO H2228 cells to a low concentration of tanespimycin (**Figure 5C**). This correlates with the cell cycle analysis showing accumulation of *SPDL1* KO H2228 cells in G2/M upon treatment with tanespimycin. We asked whether the increased expression of its subunits would result in greater activity of the CPC at kinetochores and, indeed, we detected increased levels of histone H3 phosphorylation at Ser10 only in *SPDL1* KO and not in NTC H2228 cells exposed to low concentrations of structurally distinct HSP90i for 10 d (**Figure 5D**). Importantly, we obtained similar results when EML4-ALK v3 *SPDL1* KD CUTO34 cells were exposed to a low concentration of luminespib for 10 d (**Figure 5E**).

We hypothesized that the synthetic lethal interaction between inhibition of HSP90 and loss of Spindly might be driven by unstable k-MT attachments due to increased CPC activity at kinetochores. To test this, we treated NTC and *SPDL1* KO H2228 cells with either DMSO or 12 nM tanespimycin for 12 d with or without barasertib (AZD1152-HQPA), a specific Aurora B inhibitor (45; 46) recommended by the Chemical Probes Portal (www.chemicalprobes.org/azd1152). Notably, inhibition of Aurora B with barasertib rescued the loss of cellular fitness of *SPDL1* KO H2228 cells treated with tanespimycin (**Figure 5F**). Overall, these results suggest that upon depletion of Spindly, prolonged exposure of EML4-ALK v3 NSCLC cells to low concentrations of HSP90i induces the activity of the CPC leading to spindle disruption and impairment of cellular fitness.

## Discussion

Clinical trials of HSP90i have shown therapeutic activity among EML4-ALK^+^ NSCLC patients. However, some patients have responded better than others and a satisfactory explanation for these clinical discrepancies is still lacking. Past clinical trials of HSP90i have ignored the structural differences of EML4-ALK variants despite *in vitro* studies indicating that patients harboring v1 might be more sensitive to inhibition of HSP90 (21; 22). Additionally, it has been suggested that the administration of HSP90i at or close to their maximum tolerated dose may not be the most effective therapeutic approach and the concept of using low doses of HSP90i to reduce adverse events has recently gained interest (47).

Here, we present evidence that different EML4-ALK variants do not in fact affect the sensitivity of EML4-ALK^+^ NSCLC cells to HSP90i. Although we confirmed a more rapid depletion of v1 compared to v3 upon exposure to HSP90i, EML4-ALK^+^ NSCLC cells showed decreased MAPK signaling regardless of the variant expressed and no substantial difference in sensitivity upon treatment with structurally unrelated HSP90i. Therefore, it seems unlikely that the heterogeneous responses to HSP90i observed among EML4-ALK^+^ NSCLC patients can be attributed to different EML4-ALK variants and we aimed to discover alternative mechanisms of sensitivity in EML4-ALK^+^ NSCLC cells using a systematic and unbiased approach.

CRISPR/Cas9 pooled knockout screening is a cost-effective technology for large-scale interrogations of multiple genetic perturbations simultaneously (48; 49). As chaperones are considered hubs enriched for synthetic lethal partners (25), we performed a genome-wide CRISPR/Cas9 loss-of-function screen in EML4-ALK v1 H3122 and EML4-ALK v3 H2228 cells exposed to a HSP70-inducing but non-lethal concentration of tanespimycin to investigate whether additional genetic perturbations would alter the sensitivity of EML4-ALK^+^ NSCLC cells to low concentrations of HSP90i. In the screen, we found that the knockout of *SPDL1*, the gene encoding the protein Spindly, increases the sensitivity to low concentrations of tanespimycin only in v3 H2228 and not in v1 H3122 cells. Importantly, in contrast to most EML4-ALK studies using only H3122 and H2228 cells, we validated the results of our screen in additional EML4-ALK^+^ NSCLC cell lines using several structurally distinct HSP90i and depleting Spindly with both knockout and knockdown molecular technologies. This confirmed that the findings of our screen are not restricted to a specific HSP90i chemotype or a single cell line. Beside *CDC16*, no other gene involved in the mitotic process or the formation of k-MT attachments scored significantly in our CRISPR/Cas9 screen, including the non-essential subunits of the ROD-Zwilch-ZN10 (RZZ) complex, which recruits Spindly to kinetochore (50). Moreover, we did not observe any synergistic interaction between non-lethal concentrations of tanespimycin and the microtubule-targeting agent paclitaxel. This indicates that the synthetic lethality we uncovered in EML4-ALK v3 NSCLC cells exposed to low concentrations of HSP90i specifically requires the depletion of Spindly.

Loss of Spindly did not alter the stability of EML4-ALK v3 and did not increase the sensitivity of v3 H2228 cells to tanespimycin upon treatment for only 72 h. Instead, we observed growth impairment when *SPDL1* KO EML4-ALK v3 NSCLC cells were exposed to low concentrations of HSP90i for 10 d or longer. Loss of Spindly alone had only a minimal effect on cellular fitness. Accordingly, we detected only a minor increase in the number of *SPDL1* KO cells in G2/M and a small increase in mitotic cells with uncongressed chromosomes, as well as cells with micronuclei. These observations were not surprising given the role played by Spindly in the formation of the mitotic spindle. Interestingly, we obtained similar modest effects when NTC H2228 cells were exposed to low concentrations of tanespimycin alone for 10 d. Moreover, RNA sequencing showed induced expression of genes encoding proteins involved in the formation of k-MT attachments as well as enriched positional gene sets, suggesting chromosome mis-segregation events upon treatment of NTC H2228 cells with tanespimycin alone (41). Previous studies have shown that HSP90 is involved in yeast and human kinetochore assembly (51; 52) and concentrations of tanespimycin as low as 10 nM have been reported to displace CENPH from human kinetochores (53). Moreover, higher concentrations of geldanamycin-derived HSP90i have been shown to modulate the expression of survivin in selected human cancer cell lines (54). Our data suggest that low concentrations of HSP90i modulate the formation of k-MT attachments and chromosome congression. Further research is warranted to clarify the molecular details of these events and the consequential synthetic lethal interaction with loss of Spindly.

Prolonged exposure of *SPDL1* KO H2228 cells to low concentrations of HSP90i induced a more prominent accumulation in G2/M, increased levels of prometaphase/metaphase markers and led to more severe chromosome congression abnormalities. Furthermore, we detected a greater number of interphase cells with micronuclei, suggesting more frequent chromosome mis-segregations, as well as more intact cells with fragmented nuclei. Treatment of *SPDL1* KO EML4-ALK v3 NSCLC cells with low concentrations of HSP90i for 10 d increased the expression of all subunits of the CPC as well as its activity at kinetochores. Importantly, co-treatment with a selective Aurora B inhibitor rescued the impairment of cellular fitness of *SPDL1* KO H2228 cells exposed to a low concentration of tanespimycin. We propose that loss of Spindly alone or prolonged treatment with low concentrations of HSP90i alone lead to a mild impairment of chromosome congression without any substantial effect on cellular fitness. In contrast, upon loss of Spindly, prolonged exposure of EML4-ALK v3 NSCLC cells to HSP90i increases the activity of the CPC at kinetochores leading to markedly disrupted spindles and the accumulation of cells in prometaphase. The increased activity of the CPC likely results in unstable and erroneous k-MT attachments undetected by the SAC and resulting in occasional chromosome missegregations. Cells with mis-segregated chromosomes may still undergo successful mitosis. Nevertheless, after a number of cell divisions upon increased CPC activity, impairment of chromosome congression and multiple mis-segregation events, cellular fitness would become irreparably compromised. For this reason, *SPDL1* KO cells do not show increased sensitivity to HSP90i when treated for only 72 h. The synthetic lethality we uncovered requires a prolonged exposure to low concentrations of HSP90i and the progressive accumulation of chromosome congression defects (**Figure 6**).

**Figure 6.**
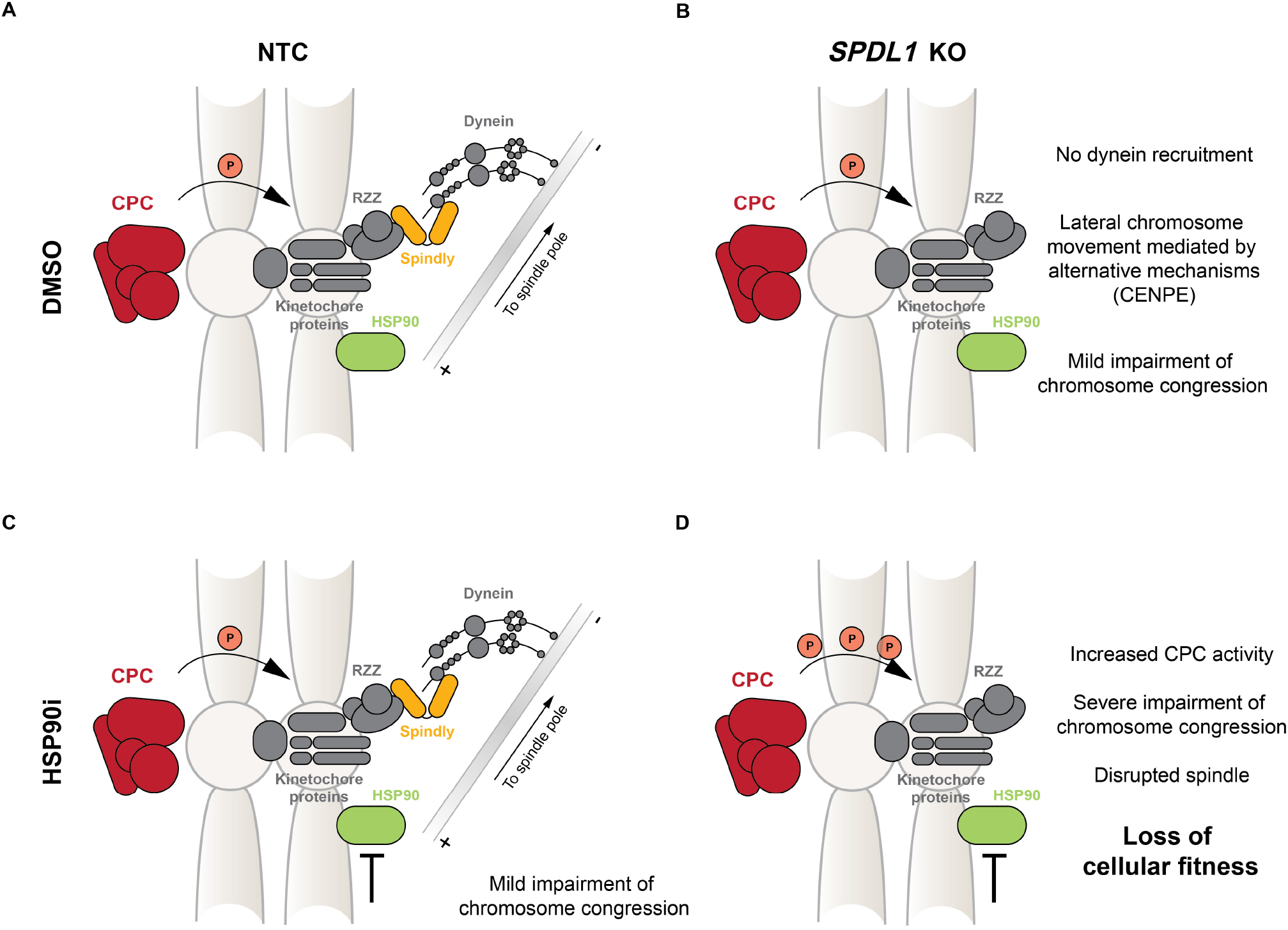
Diagrammatic model showing how loss of Spindly impairs the fitness of EML4-ALK v3 NSCLC cells exposed to low concentrations of HSP90i. (**A**) In the absence of external perturbations Spindly interacts with the RZZ complex to recruit dynein to kinetochores and facilitate chromosome congression. (**B**) Upon loss of Spindly, chromosomes can still congress to the spindle equator via dynein-independent mechanisms. However, this leads to a mild impairment of chromosome congression and occasional mis-segregations with no prominent impact on the fitness of EML4-ALK v3 NSCLC cells. (**C**) Prolonged treatment with low concentrations of HSP90i does not decrease the fitness of EML4-ALK v3 NSCLC cells but induces a mild impairment of chromosome congression as HSP90 participates in kinetochore assembly. (**D**) In the absence of Spindly, prolonged exposure of EML4-ALK v3 cells to low concentrations of HSP90i induces the activity of the CPC at kinetochores and leads to mitotic delay, severe impairment of chromosome congression, spindle disruption, chromosome mis-segregations and loss of cellular fitness.

We observed loss of cellular fitness upon combined depletion of Spindly and prolonged exposure to low concentrations of HSP90i exclusively in EML4-ALK v3, and not in v1, NSCLC cells. It has been shown that while both v1 and v3 localize to cytoplasmic granules, only v3 binds to microtubules (5; 6; 7). It is tempting to speculate that v3 may bring the tyrosine kinase activity of ALK to the spindle where the phosphorylation of proteins involved in chromosome congression might generate a specific vulnerability to loss of Spindly and to prolonged treatment with low concentrations of HSP90i. Alternatively, the mitosis-retained microtubule-stabilizing activity of EML4-ALK v3 could contribute to selective dependencies (55). Further investigations on the differences between EML4-ALK v1 and v3 and their impact on specific synthetic lethal interactions are warranted.

Most studies on EML4-ALK^+^ NSCLC are carried out using only v1 H3122 and v3 H2228 patient-derived cell lines or artificial overexpression models. To corroborate our data, we sourced two additional EML4-ALK^+^ NSCLC patient-derived cell lines and reproduced our original findings. Our results suggest that EML4-ALK v3 NSCLC patients with reduced levels of Spindly or loss-of-function mutations in the *SPDL1* gene could potentially show more favorable responses to low doses of HSP90i. Interestingly, lung adenocarcinoma ranks third by genetic alteration frequency in the *SPDL1* gene according to cancer genomics data available in cBioPortal (56) (**Figure S6A**). Mutations in the *SPDL1* gene have not been annotated yet but many of those found in the genome of cancer patients are predicted using canSAR (57) to be moderately or highly disruptive. We did not retrieve any overlap between ALK fusions and mutations in the *SPDL1* gene in lung adenocarcinoma. However, the *SPDL1* gene was not profiled in the vast majority of these patient samples (**Figure S6B**). We encourage retrospective studies on genomics data if these were collected during clinical trials of HSP90i enrolling EML4-ALK^+^ NSCLC patients. Together with our results, these analyses could lead to a better understanding of the heterogeneous responses to HSP90i, foster better stratification of EML4-ALK^+^ NSCLC patients in future clinical trials and establish Spindly as a potential sensitivity biomarker.

In summary, we showed that, despite the more rapid depletion of v1 upon inhibition of HSP90, the presence of EML4-ALK v1 or EML4-ALK v3 per se does not determine the sensitivity of NSCLC cells to structurally distinct HSP90i. Using an unbiased and large-scale CRISPR/Cas9 screening approach we investigated the effect of thousands of genetic perturbations on the sensitivity of EML4-ALK^+^ NSCLC cells to low concentrations of HSP90i in what is, to our knowledge, the first EML4-ALK-focused synthetic lethal screen. We found that upon loss of Spindly EML4-ALK v3, but not v1, NSCLC cells become more sensitive to prolonged treatment with concentrations of HSP90i that are much lower than those routinely used *in vitro* and in clinical studies to achieve HSP90 protein client depletion. Such low concentrations of HSP90i are less likely to induce the adverse events commonly associated with the administration of these drugs to patients (18; 19).

## Methods

### Cell lines

NCI-H2228 [H2228] (ATCC, CRL-5935, female, human) and NCI-H3122 [H3122] (NCI DTP Developmental Therapeutics Program, male, human) cells were cultured at 37°C in RPMI medium (Sigma-Aldrich) supplemented with 10% FBS (PAN Biotech), were validated by STR profiling at the Institute of Cancer Research and were tested for mycoplasma infection. CUTO8 (*EML4-ALK* E13;A20, v1) cells were derived from a pleural based mass of a female patient who was age 66 at diagnosis with lung adenocarcinoma and while progressing on crizotinib (other prior therapies included carboplatin/docetaxel/bevacizumab). CUTO34 (*EML4-ALK* E6;A20, v3) were derived from a pleural based mass of a male patient who was age 46 at diagnosis with lung adenocarcinoma while progressing on alectinib (no other prior therapies). Both cell lines were obtained from the laboratory of Robert Doebele, were cultured at 37°C in RPMI medium (Sigma-Aldrich) supplemented with 10% FBS (PAN Biotech), were submitted to ATCC for STR profiling, were confirmed to be human but did not match any profile in the ATCC or DSMZ STR database and were tested for mycoplasma infection.

### Western blotting

Cells were lysed in radioimmunoprecipitation assay (RIPA) buffer supplemented with a cocktail of protease (Roche) and phosphatase inhibitors (Sigma-Aldrich). Protein concentration was determined with the Pierce BCA Protein Assay Kit (Thermo Fisher Scientific). Lysates were resolved by SDS-PAGE and transferred to nitrocellulose membranes using the iBlot2 Gel Transfer Device (Thermo Fisher Scientific). ALK (3633), p44/p42 MAPK Erk1/2 (4695), phospho-p44/p42 MAPK Erk1/2 Thr202/Tyr204 (4370), c-PARP (9541), NQO1 (3187), cyclin B1 (4135), securin (13445), histone H3 (9715), phospho-histone H3 Ser10 (9706) and vinculin (13901) antibodies were purchased from Cell Signaling Technology. GAPDH antibody (MAB374) and Cas9 antibody (MAC133) were purchased from Millipore. HSP72 (ADI-SPA-810) antibody was purchased from Enzo Life Sciences. Spindly antibody (CSB-PA842662LA01HU) was purchased from Cusabio.

### Cellular fitness measurements

For concentration-response curves, 4,000 cells/well were seeded in 96-well plates and exposed after 24 h to either DMSO or drugs as indicated. After 72 h, cell viability was measured using CellTiter-Glo (Promega). For growth curves, 100,000 cells/well were seeded in 6-well plates and exposed after 24 h to either DMSO or 12 nM tanespimycin. After 72 h, cells were counted, reseeded and treated again to continue the experiment for up to 24 d. For cell viability measurements in 6-well or 12-well plates, cells were treated as indicated then washed with cold PBS, fixed with 3.7% formaldehyde in PBS for at least 30 min and then stained with crystal violet solution (20% methanol) for at least 30 min. For quantification, the crystal violet staining was extracted using a 50% ethanol solution and absorbance read at 590 nm.

### Cas9 activity assay

Cas9-expressing H3122 and H2228 cells were transduced with the pXPR-011-sgEGFP construct, which allows for the expression of EGFP and an sgRNA targeting the *EGFP* gene, and selected with puromycin. After 8 d, cells were analyzed by flow cytometry on a BD LSR II (BD Biosciences). Cas9 endonuclease was considered sufficiently active when EGFP^+^ cells amounted to less than 25%.

### Expansion of the Brunello library

The Brunello library (28) was purchased from Addgene (73178). In all, 100 ng of library was added to 25 µL of Endura electrocompetent cells (Lucigen). Cells were electroporated using the MicroPulser (BioRad, Ec1 setting), immediately recovered in 975 µL of pre-warmed recovery medium, transferred to a tube containing an additional 1 mL of recovery medium and shaken at 250 rpm for 1 h at 37°C. We performed a total of 4 electroporations on 4 independent aliquots of competent cells, which were then pooled, plated on 4 pre-warmed 24.5 cm^2^ bioassay ampicillin agar plates and grown for 14 h at 32°C. Colonies were harvested using a cell scraper and plasmids were purified using the HiSpeed Plasmid Maxi Kit (QIAGEN). We confirmed homogeneous library representation by next-generation sequencing (**Figure S2C**).

### Genome-wide CRISPR/Cas9 pooled screen

In all, enough cells were seeded and kept in culture throughout the entire screen to ensure a 500x coverage of the Brunello library. After 24 h, cells were transduced with the Brunello lentiviral library at a multiplicity of infection of around 0.3-0.5. Puromycin selection was initiated after 24 h. After 5 d, selected cells were split in two branches and treated with either DMSO or 12 nM tanespimycin for 12 d. At the end of the screen, genomic DNA was extracted with the Blood and Cell Culture DNA Maxi Kit (QIAGEN). A nested PCR was performed to amplify and barcode the sgRNA cassettes out of a sufficient amount of genomic DNA to ensure a 500x coverage of the Brunello library. The PCR was performed using Herculase II Fusion DNA Polymerase (Agilent Technologies) and primers are listed in **Table S5**. Barcoded PCR amplicons from different samples were quantified on an agarose gel, pooled in equimolar amounts, purified using the QIAquick Gel Extraction Kit (QIAGEN) and sequenced on a NextSeq 500 in High Output mode and using a 75-bp single-end protocol (Source BioScience, UK). For each condition, 2 independent biological replicates were performed.

### Reverse transcription quantitative polymerase chain reaction

RNA was extracted using the RNeasy Mini Kit (QIAGEN) and cDNA was produced using the High-Capacity cDNA Reverse Transcription Kit (Thermo Fisher Scientific). The quantitative polymerase chain reaction was performed using the Power SYBR Green PCR Master Mix (Thermo Fisher Scientific) on a ViiA 7 Real-Time PCR System (Thermo Fisher Scientific).

### Cell cycle analysis

Cells were harvested, washed and resuspended in PBS, fixed with cold 70% ethanol for 15 min, centrifuged and resuspended in PBS to rehydrate for 10 min. Cells were then incubated at 37°C for 30 min after the addition of RNase A (QIAGEN) to a final concentration of 100 µg/mL. Propidium iodide (Sigma-Aldrich) was added to a final concentration of 50 µg/mL and cells were analyzed on a BD LSR II (BD Biosciences) flow cytometer.

### Immunofluorescence

Cells were grown on coverslips, washed with PBS, fixed with methanol at -20°C for 15 min, washed with PBS, incubated with blocking buffer (3% BSA, 0.1% Triton X-100 in PBS) for 15 min at room temperature, incubated with primary antibodies in blocking buffer for 30 min at room temperature, washed 3 times with PBS, incubated with secondary antibodies in blocking buffer for 30 min at room temperature in the dark, washed 3 times with PBS and once with distilled water, mounted on slides using ProLong Diamond Antifade Mountant with DAPI (Thermo Fisher Scientific) and left to dry over night at room temperature. β-tubulin (T4026) and γ-tubulin (T3559) were purchased from Sigma-Aldrich. Alexa Fluor 488 donkey anti-mouse IgG (A21202) and Alexa Fluor 546 donkey anti-rabbit IgG (A10040) antibodies were purchased from Thermo Fisher Scientific. Images were taken using a Zeiss LSM 700 inverted confocal microscope with a Plan-Apochromat 40x/1.3 Oil DIC M27 objective or a Metafer4 automated slide scanner including a Zeiss Axio Imager.Z2 upright microscope with an EC Plan-Neofluar 40x/0.75 dry M27 objective. Images were edited using the Zen 2009 and VSViewer software.

### Gene expression profiling

Cells were lysed in Nucleic Acid Purification Lysis Solution (Thermo Fisher Scientific) and RNA was extracted using the MagNA Pure 96 (Roche). Samples were quantified by Nanodrop and sequenced on a BGISEQ-500 using a 100-bp paired-end protocol (BGI, Hong Kong). Reads were aligned to the human genome (GRCh38.p13) using STAR (2-pass) and differential gene expression analysis was performed using DESeq2. For each condition, 4 independent biological replicates were performed. RNA-seq data will be available in Gene Expression Omnibus (GEO).

### Histone extraction

Cells were washed in cold PBS and resuspended in 0.5% v/v Triton X-100 in PBS supplemented with a cocktail of protease (Roche) and phosphatase inhibitors (Sigma-Aldrich) at a cell density of 10^7^ cells/mL. After 10 min on ice, samples were centrifuged at 6,500*g for 10 min at 4°C. Cells were washed with half volume of 0.5% v/v Triton X-100 in PBS and centrifuged again as before. Pellets were resuspended in 0.2 M HCl at a cell density of 4×10^7^ cells/mL to extract histones over night at 4°C. Samples were centrifuged at 6,500*g for 10 min at 4°C and the supernatants were neutralized using 2 M NaOH at 1/10 of the volume.

### Quantification and statistical analysis

PoolQ v1.0.0 and STARS v1.2 were used to analyze data from the CRISPR/Cas9 screen and calculate Average Scores and False Discovery Rate (FDR). In line with convention, an arbitrary threshold of FDR<0.1 was chosen to select significant screening hits (FDR<0.01 for high-confidence hits). DESeq2 was used to analyze RNA-Seq data and calculate fold changes in gene expression between samples as well as FDR. In line with convention, arbitrary thresholds of fold change>2 and FDR<0.01 were chosen to select the most significant gene expression changes. GSEA v4.2.3 was used to perform gene set enrichment analysis with default settings (1,000 permutations). Only gene sets enriched with FDR<0.05 were considered. Error bars in the figures were calculated using GraphPad Prism and always refer to s.d. of biological replicates. Additional statistical details, including number of biological replicates, are indicated in the figure legends.

## Acknowledgements

The authors thank Steven Whittaker (Institute of Cancer Research, ICR) for help and useful discussion on the CRISPR screen, Louise Howell (ICR) for help with confocal microscopy and Ian Titley (ICR) for help with flow cytometry. We thank Jon Pines (ICR) for helpful comments. P. Workman acknowledges program grant support from Cancer Research UK (CRUK grant numbers C309/A11566 and C2739/A22897) and support from the ICR (London, United Kingdom). P. Workman acknowledges additional grant support from the Wellcome Trust (212969/Z/18/Z), Cancer Research UK (C35696/A23187), Mark Foundation and Chordoma Foundation and he is a Cancer Research UK Life Fellow. ICR authors acknowledge infrastructure support from the CRUK Centre at the ICR.

## Author Contributions

M.P.L. designed and performed the CRISPR/Cas9 screen and all experiments. C.Z. performed reads alignment and DESeq2 analyses for the gene expression profiling data. A.T.L. and R.C.D. provided additional cell lines and support with experiments. M.P.L., P.C.A. and P.W. conceived the study and wrote the manuscript. All authors approved the manuscript.

## Manuscript preparation

This manuscript was prepared in Overleaf (http://www.overleaf.com) using a modified version of the RoyleLab bioRχiv template.

## Supplementary Information

### List of Supplementary Tables

**Table S1**. Fitness genes depleted in the DMSO T_12_ sample versus the T_0_ sample of the H3122 and H2228 CRISPR/Cas9 screens.

**Table S2**. Genes depleted or enriched in the tanespimycin T_12_ sample versus the DMSO T_12_ sample of the H3122 and H2228 CRISPR/Cas9 screens.

**Table S3**. Gene expression profiling fold changes.

**Table S4**. Gene sets enriched according to GSEA.

**Table S5**. Primers used.

**Figure S1.**
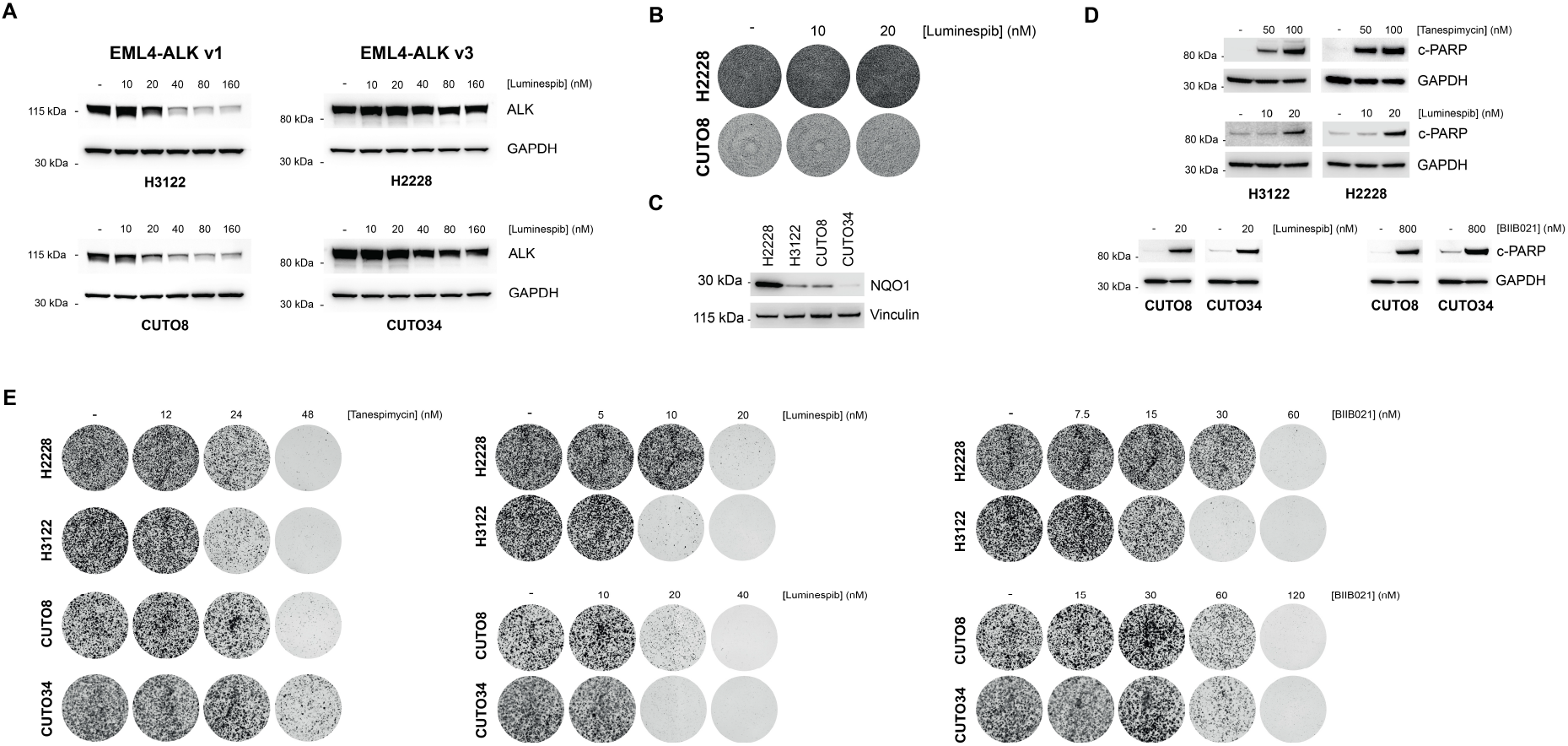
Treatment of EML4-ALK^+^ NSCLC cells with HSP90i. **(A)** Western blotting of EML4-ALK^+^ NSCLC cells exposed to luminespib at the indicated concentrations for 24 h. GAPDH was used as a loading control. **(B)** Crystal violet staining of H2228 and CUT08 cells exposed to luminespib at the indicated concentrations for 24 h. **(C)** Western blotting of EML4-ALK^+^ NSCLC cells. Vinculin was used as a loading control. **(D)** Western blotting of EML4-ALK^+^ NSCLC cells exposed to HSP90i at the indicated concentrations for 72 h. GAPDH was used as a loading control. **(E)** Colony forming assay of EML4-ALK^+^ NSCLC cells exposed to HSP90i at the indicated concentrations for 12 d.

**Figure S2.**
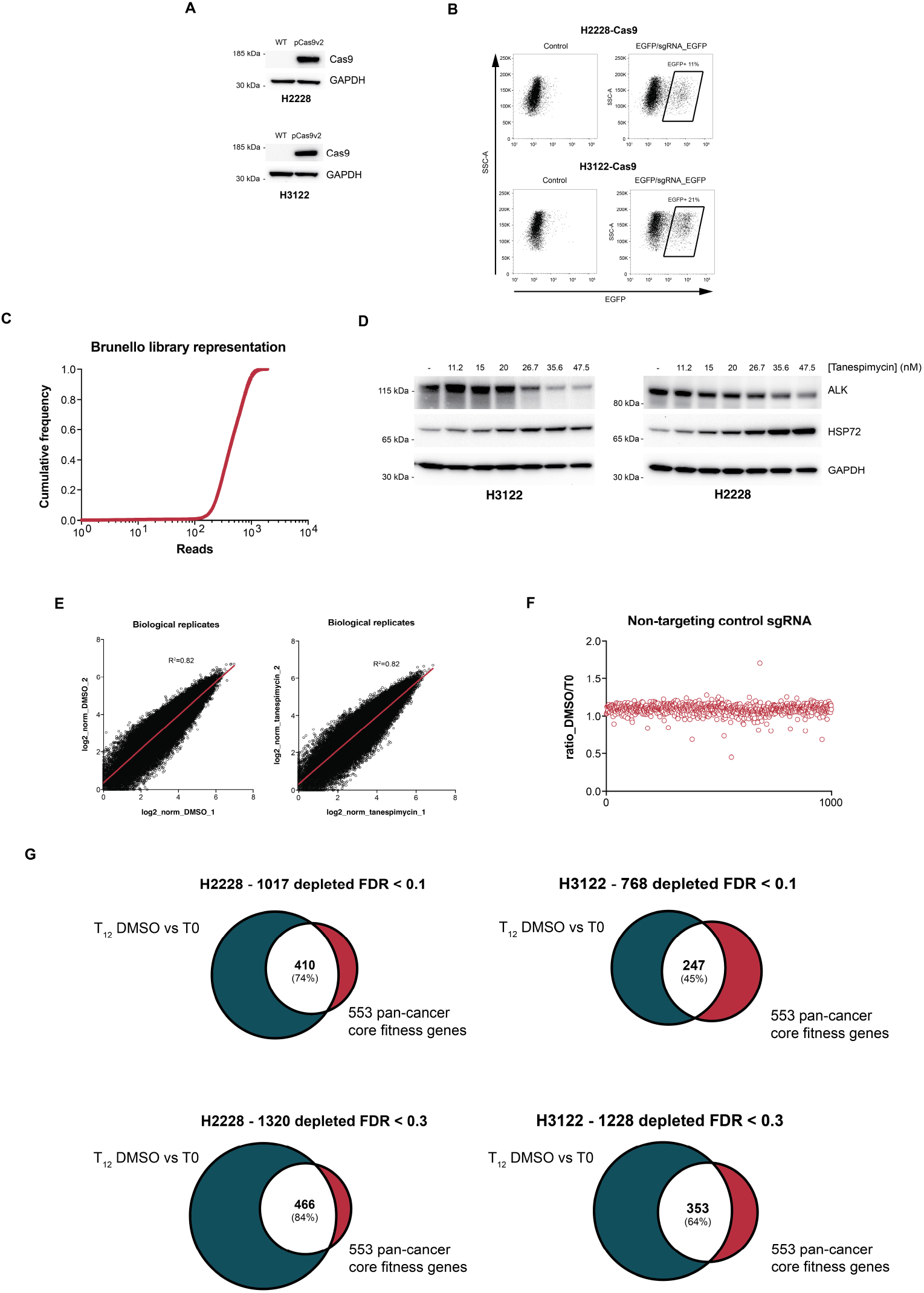
Genome-wide CRISPR/Cas9 knockout screen. **(A)** Western blotting of H2228 and H3122 cells transduced with pLEX_311-Cas9v2. GAPDH was used as a loading control. **(B)** Flow cytometry analysis of Cas9-expressing H2228 and H3122 cells transduced with the EGFP/sgRNA_EGFP vector. EGFP^+^ cells<25% indicate ideal Cas9 activity. **(C)** Cumulative frequency plot showing homogeneous representation of sgRNAs in the amplified Brunello library. **(D)** Western blotting of H3122 and H2228 cells exposed to the indicated concentrations of tanespimycin for 12 d. GAPDH was used as a loading control. **(E)** Correlation plots of normalized reads of biological replicates from the H2228 CRISPR/Cas9 screen. **(F)** Ratio of normalized reads of Brunello 1,000 NTC sgRNAs between the T_12_ DMSO sample and the T_0_ sample. **(G)** Venn diagrams showing pan-cancer core fitness genes depleted in the T_12_ DMSO sample versus the T_0_ sample of the H2228 and H3122 CRISPR/Cas9 screens.

**Figure S3.**
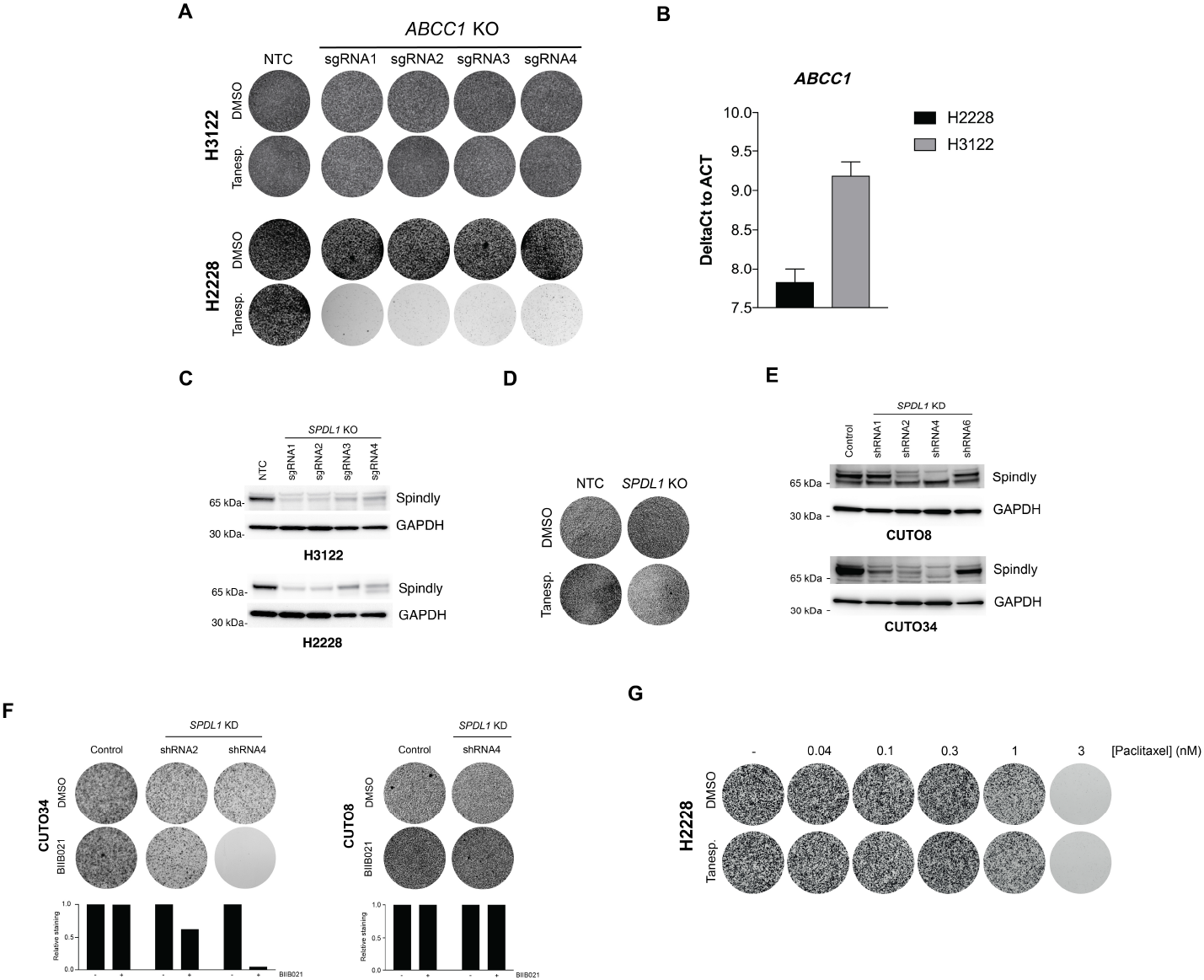
CRISPR/Cas9 screen hit validation experiments. **(A)** Crystal violet staining of NTC and *ABCC1* KO H3122 and H2228 cells exposed to DMSO or 12 nM tanespimycin for 12 d. **(B)** RT-qPCR analysis of *ABCC1* expression in H2228 and H3122 cells. The higher the Ct (Cycle threshold) value the lower the expression. Data are normalized to actin expression. Error bars, s.d. of 3 biological replicates. **(C)** Western blotting of Cas9-expressing H3122 and H2228 cells transduced with a NTC sgRNAor with sgRNAs targeting *SPDL1*. GAPDH was used as a loading control. **(D)** Crystal violet staining of NTC and *SPDL1* KO CUTO8 cells exposed to DMSO or 20 nM tanespimycin for 12 d. **(E)** Western blotting of CUTO8 and CUTO34 cells transduced with a control shRNA or with shRNAs targeting *SPDL1*. GAPDH was used as a loading control. **(F)** Crystal violet staining and quantification of control and *SPDL1* KD CUTO34 and CUTO8 cells exposed to DMSO or 30 nM BIIB021 for 12 d. **(G)** Colony forming assay of H2228 cells exposed to paclitaxel at the indicated concentrations and to either DMSO or 12 nM tanespimycin for 12 d.

**Figure S4.**
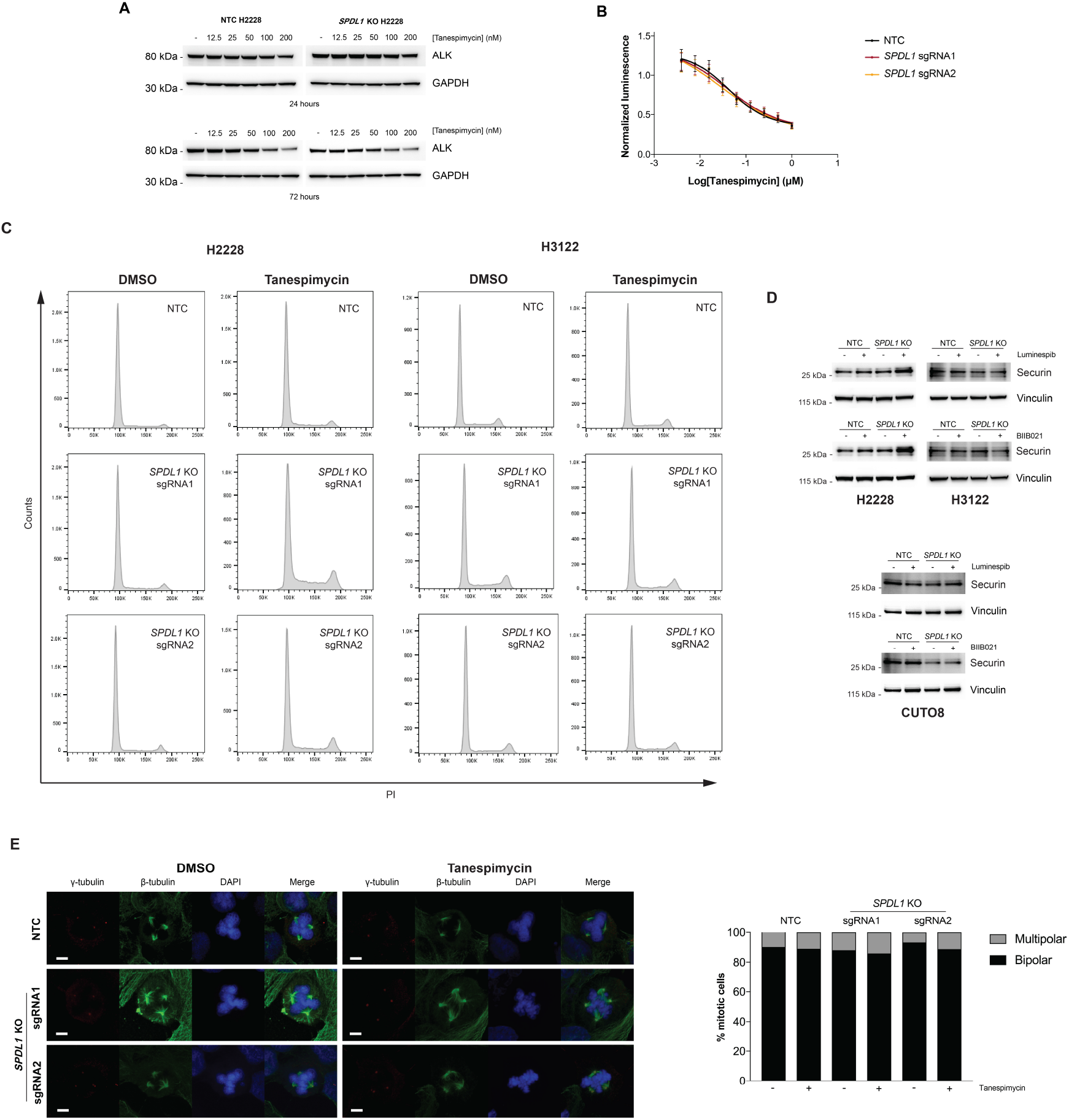
Stability of EML4-ALK v3, cell cycle and immunofluorescence analysis of EML4-ALK^+^ NSCLC cells upon loss of Spindly and treatment with HSP90i. **(A)** Western blotting of NTC and *SPDL1* KO H2228 cells exposed to tanespimycin at the indicated concentrations for 24 or 72 h. GAPDH was used as a loading control. **(B)** Concentration-response curves of NTC and *SPDL1* KO H2228 cells exposed to tanespimycin as indicated for 72 h. Data are normalized to vehicle-treated cells set to 100. Error bars, s.d. of 6 biological replicates. **(C)** Cell cycle analysis of NTC and *SPDL1* KO H2228 (left) and H3122 (right) cells exposed to DMSO or 6 nM tanespimycin for 10 d. Percentage of cells are reported in Figure 3B. **(D)** Western blotting of NTC and *SPDL1* KO H2228 and H3122 cells exposed to DMSO or 3 nM luminespib or 10 nM BIIB021 and NTC and *SPDL1* KO CUTO8 cells exposed to DMSO or 3 nM luminespib or 15 nM BIIB021 for 10 d. **(E)** Immunofluorescence analysis and quantification of NTC and *SPDL1* KO H2228 cells treated as in (C). More than 200 mitotic cells per condition were counted to determine percentages. Scale bar, 5 μm.

**Figure S5.**
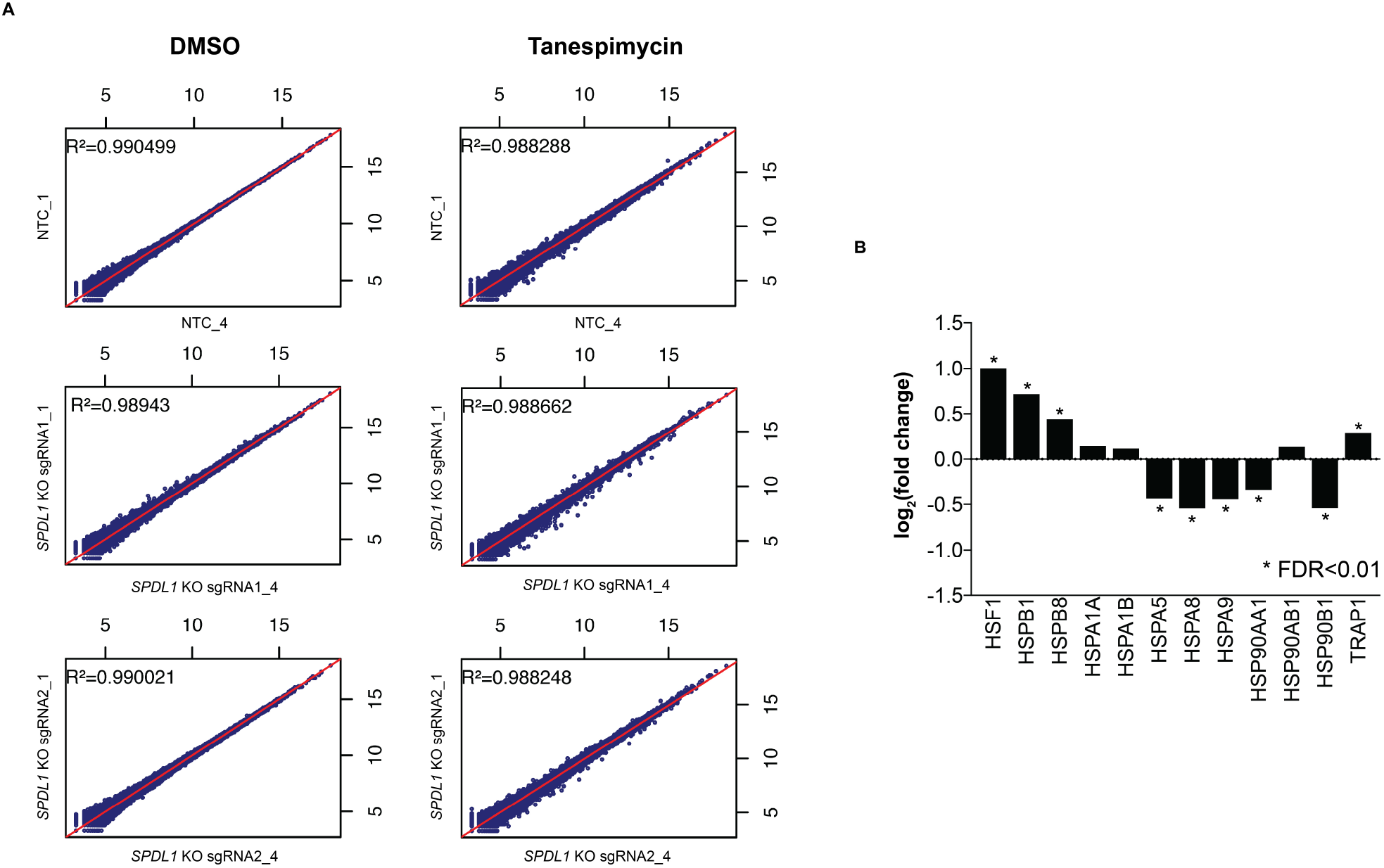
Gene expression profiling of H2228 cells upon loss of Spindly and treatment with tanespimycin. **(A)** Representative correlation plots of biological replicates from the RNAseq gene expression profiling. **(B)** Gene expression changes in the heat shock response upon exposure of H2228 cells to 6 nM tanespimycin for 10 d. The Iog2 of the fold change to control (H2228 cells exposed to DMSO) is shown. Expression changes with a FDR<0.01 are indicated with an asterisk.

**Figure S6.**
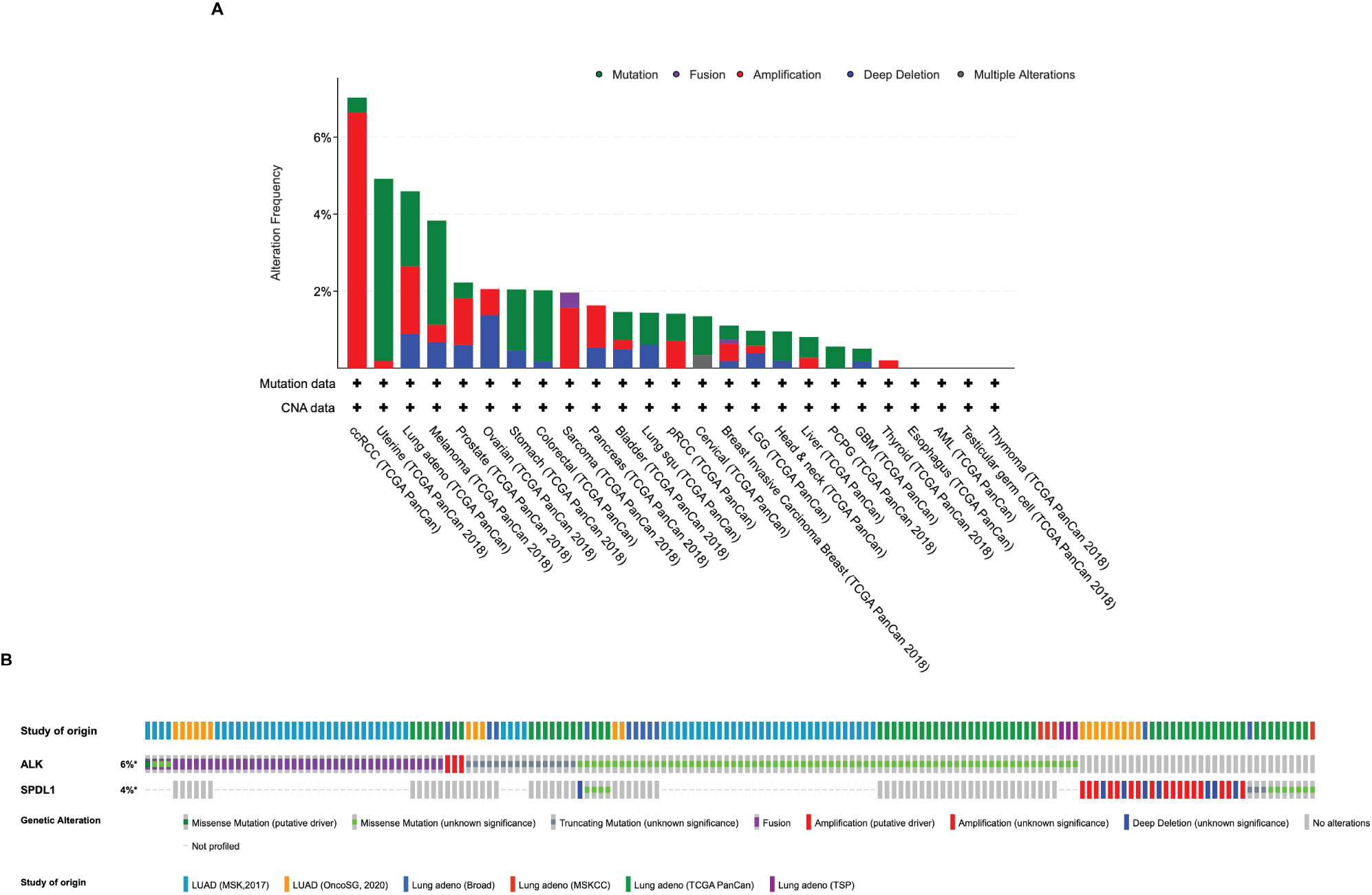
Cancer genomics data from cBioPortal. (**A**) Cancer types raked by alteration frequency in the *SPDL1* gene. Only studies with at least 100 cases were considered. (**B**) Oncoprint showing genetic in the *ALK* and *SPDL1* genes in lung adenocarcinoma.

